# Does metal pollution affect stoichiometry of soil-litter food webs?

**DOI:** 10.1101/2019.12.17.879205

**Authors:** Michel Asselman, Łukasz Sobczyk, January Weiner, Stefan Scheu, Anna Rożen

## Abstract

To date the field of ecological stoichiometry has focused mainly on aquatic systems concentrating on macro-elements. We investigated terrestrial systems and included microelements to study the elemental transfer in the detritivorous food web. We compared food webs of six sites differing in the type and degree of metal pollution along two forest transects contaminated with copper or zinc. We measured 11 elements in litter, herbivores, detritivores, predators and omnivores. Based on elemental concentrations of elements differences between trophic groups were visualized using PCA. At all sites litter C:N, C:P, C:K and C:Na ratios were higher than in animals. Invertebrate trophic groups were significantly different from each other in C:Cu, C:Zn and C:Ca ratios. The calculated resource:consumer N:P ratio suggests that invertebrates in studied forests are N limited and not P limited. Similar patterns at all sites suggests that metal pollution at the studied intensity slightly affects the transfer of elements in the terrestrial macro-invertebrate food web.

## Introduction

Since the work of Lindeman (1942) the flow of energy through food webs has been an important aspect of ecology. Energy is counted only in Joules, hence Hessen et al. (2004) suggested that it would be more convenient to use carbon (C) as a currency for the flow of both energy and matter at the same time. Further C can be measured together with other biological key elements such as e.g. N, P, Fe, Zn, Cu, K, Na. Among other elements crucial for organism functioning are those which are necessary to build tissues of organisms like nitrogen and phosphorus, and other which are constituents of enzymes (e.g. Fe, Zn, Cu, Mn, Ca, Mg) or are involved in other processes (e.g. K, Na). When the total amount of elements has been measured within different compartments of an ecosystem, the relative abundance of these elements can be used as determinants of ecosystem processes. The field of ecology dealing with these relative abundances of elements is called “ecological stoichiometry” (Sterner and Elser, 2002).

Stoichiometric differences between different groups of invertebrates are relatively small, compared to the differences between animals and plants (Cease and Elser, 2013). This could result in the largest elemental differences at the first trophic link, from plant/detritus to herbivores/detritivores and a lesser one between herbivores and predators (Bradshaw et al., 2012). Even though there have been great advances in the field of ecological stoichiometry, much still need to be unfolded. Studies until today mainly focused on macro-elements (C, N, P), and were conducted mostly in aquatic environments. Studies on multi-elemental and multi species/trophic level such as Karimi and Folt (2006), Bradshaw et al. (2012), Filipiak and Weiner (2014), Filipiak et al. (2016) are scarce.

Essential metals such as copper and zinc have been mainly studied as pollutant and but received little attention from a nutritional point of view as microelements stoichiometrically interacting with other macro- and microelements. Zn and Cu are important key constituents of enzymes and proteins, however, they were taken into account first of all in the field of stress ecology and ecotoxicology because of the negative effects of high doses of these elements on living organisms (Tyler 1984). Literature data show that growth of individuals is negatively affected by metals (e.g. Donker et al.,1993, Rożen, 2006). Body composition and stoichiometry are related to ontogeny (Boros et al., 2015) what suggest possible impact of metals on body stoichiometry of invertebrates.

The interaction between elements has been studied in regard to the uptake by plants (Siedlecka, 1995), revealing that changes in availability of each group can affect the uptake of the other (Lin and Wu, 1994; Liu et al., 2003; Chen et al., 2007; Peng et al, 2008). In the detritivorous food web, litter decay processes and metal accumulation in invertebrates have so far mainly been linked to soil type, soil metal concentration and dominant tree species, including their effect on chemical composition of leaf litter (Vesterdal, 1999; Sariyildiz et al., 2005).

It has been shown in the previous studies on metal pollution transect in Olkusz forest that the metal pollution has negative effects upon litter soil invertebrates: their density (Enchytraeids – Tosza et al. 2010), sensitivity to additional stressor (Carabid beetles – Stone et al., 2001, Łagisz and Laskowski, 2007), however, a positive correlation has also been found between metal concentration and body mass of beetles (Zygmunt et al., 2007). The studies concerning the effect of pollution on microbial communities in the above mentioned transects did not bring any clear answer (Stefanowicz et al., 2007, Chodak et al., 2013). The response of invertebrates to heavy metals (accumulation in the body) depends on various factors: habitat, diet, physiological response and therefore varioius taxonomic groups differ in the ability to accumulate and eliminate metals (Gall et al., 2015).

Our hypothesis was that the costs of detoxification (e.g. production of metallothioneins, storage of metals in granules, increased release of metals in excess) in heavy polluted sites will cause higher energetic costs (decrease of fat reserves) resulting in changes in concentrations of other elements, especially the ones engaged in energetic processes (Mn, Fe, P), and will affect the ratios between elements e.g. C:N ratio or N:P ratio. Our question is whether any differences between trophic groups exist in body composition and stoichiometry due to metal pollution.

## Methods and Materials

Two metal polluted areas were chosen on the base of previous studies: the zinc polluted Olkusz region in southern Poland and the copper polluted Legnica region in Western Poland (Niklińska et al., 2006).

In the both polluted areas transects of sites differing in metal pollution have been established (from heavy polluted to reference). Especially intensively studied was the Olkusz Forest, yielding numerous papers concerning litter decomposition (Niklińska et al., 2005; Niklińska et al., 2006), litter fauna (Tosza et al., 2010), microorganisms (Niklińska et al., 2005; Niklińska et al., 2006; Chodak et al., 2013). Similar data on Głogów Forest are available as well (Niklińska et al., 2006; Stefanowicz et al., 2008; Chodak et al., 2013).

The Olkusz region received large inputs of zinc since the medieval period due to ore mining activities and since the 1960s from two large zinc smelters, which produced ca. 118 t × m^−2^ of smelter dust. It causes that the Zinc concentrations in soil/litter locally exceed 4600 mg × kg^−1^ (Niklińska et al., 2005; Niklińska et al., 2006). On the other hand, the Głogów area is the major copper producing center in Poland. There are two copper smelters and four copper ore mines in the area causing copper concentrations in soil/litter up to 1200 mg × kg^−1^ (Niklińska et al., 2006).

In the forests of the both areas the main tree species is *Pinus sylvestris*, admixed with a small number of other tree species (*Quercus* sp. and *Betula* sp.). The soils are sandy, podzolized and acidic. The sites near Olkusz have well developed mor humus layers (5 cm) which are much thinner at the sites near Głogów (1-2 cm). In the Transects with hree sites each were determined in each of the both regions (Olkusz and Głogów) to represent various levels of pollution (heavy polluted -H, moderately polluted – M and reference -R). The reference sites in both transects were established in similar forest types, distant from pollution sources, with a background concentration of heavy metals. Coordinates of the sites and their distance to the smelters are given in Table 1.

**Table 1:** Geographical location of the studied pollution transects (Głogów and Olkusz), their distance to the smelter, concentrations of copper and zinc in litter. GH – Głogów heavily polluted, GM – Głogów moderately polluted, GR – Głogów reference site; OH – Olkusz heavily polluted, OM – Olkusz moderately polluted, OR – Olkusz reference site

| Site | Lat.<br>Degr.<br>Nord | Long.<br>Degr.<br>East | Distance<br>from<br>smelter<br>(km) | Zinc (mg/kg) ± SD | | | Copper (mg/kg) ± SD | | |
| --- | --- | --- | --- | --- | --- | --- | --- | --- | --- |
|  |  |  |  | Litter |  |  | Litter |  |  |
| GH | 51°44' | 16°1' | 5.8 | 70.4<sup>A</sup> | ± | 4.8 | 77.1<sup>A</sup> | ± | 32.5 |
| GM | 51°44' | 16°5' | 8.6 | 56.6<sup>B</sup> | ± | 15.6 | 24.8<sup>B</sup> | ± | 16.2 |
| GR | 51°47' | 16°1" | 11.2 | 66.4<sup>AB</sup> | ± | 10.0 | 13.9<sup>B</sup> | ± | 18.4 |
| OH | 50°17' | 19°29' | 2.5 | 449<sup>a</sup> | ± | 15.3 | 20.4<sup>ab</sup> | ± | 7.8 |
| OM | 50°19' | 19°30' | 3.9 | 158<sup>b</sup> | ± | 19.2 | 55.5<sup>b</sup> | ± | 33.8 |
| OR | 50°32' | 19°38' | 31.9 | 147<sup>b</sup> | ± | 2.9 | 8.2<sup>a</sup> | ± | 1.1 |

Statistically significant differences between Głogów sites are marked with capital letters, and between Olkusz sites with lower case letters

The climate of both areas is temperate, with mean annual temperature 8.0°C (Olkusz) and 8.9°C (Głogów), and annual average precipitation 600-700 mm (Olkusz) and 500-550mm (Głogów) (Chodak et al., 2013).

The litter layer was sampled at four locations per site in June 2011 at the Olkusz sites (OH – Olkusz heavy polluted, OM – Olkusz moderately polluted, OR – Olkusz reference site) and June 2012 at the Głogów sites (GH – Głogów heavy polluted, GM – Głogów moderately polluted, GR – Głogów reference site). Litter was collected from the forest floor, mixing freshly fallen and partially decomposed litter. I the laboratory samples were dried using a vacuum drier at 50 °C for 48 h, ground and stored frozen at −20°C in an airtight container until further use.

To collect macro invertebrates, 500 pitfall traps were placed at each site in the soil at 1 m from one another. Traps consisted of two 200 ml plastic cups one in another, filled with 100 ml of 70 % ethanol (EtOH). A 10 cm diameter lid was placed ±2 cm above each trap to prevent dilution by rainfall. Invertebrates were collected from the traps daily, and EtOH was replaced every other day for a period of 14 days during June 2011 in Olkusz and June 2012 in Głogów. Ethanol was chosen as a trapping liquid in pitfall traps as it was shown not to have any significant effects on invertebrates’ body stoichiometry during short time (3 days) exposition (Rożen et al., 2015).

In the laboratory, animals were sorted to the lowest taxonomic level (species or morphospecies). In the present study only individuals with specified taxonomic and trophic position have been included, provided that sufficient material for analysis of multiple samples per group (see supplement 1 for number of animals per taxonomic level used) was available. Animals were rinsed with deionized water to eliminate dust on the body and dried using a lyophilizer (Christ BETA2-8 LDplus, Martin Christ Getrieftrocknunganslagen GmbH, Germany) at −30 °C (37 Pa) for 24 h and −76 °C (0. 1 Pa) for 12 h and then stored at −20 °C in airtight containers until further use.

Prior to analyses, samples of animals and litter were homogenized and lyophilized at −30 °C for 24 h (37 Pa) once more to eliminate any moisture taken up from the atmosphere during the process of homogenization. C and N contents were examined using a CHNS analyzer (Vario EL III Elemental Analyzer, Elementar Analysensysteme GmbH, Germany). For other elements (Na, Mg, P, K, Ca, Mn, Fe, Cu, Zn) samples were prepared using digestion bombs (Heinrichs et al., 1986). Approximately 100 mg of animal or litter sample was digested using 2 ml 65 % Suprapur^®^ nitric acid (Sigma-Aldrich) in Teflon containers and pressure digested for 9 h in 185 °C. Samples were filtered and rinsed with deionized water into 50 ml volumetric flasks. Elemental concentrations were then measured using an Inductively Coupled Plasma Analyzer (Optima 5300DV ICP-OES, Perkin Elmer, Rodgau, Germany). Measurements were recalculated to milligrams per kilogram dry weight. As a reference material we used sulfanilic acid for C and N analysis, and Certified Reference Materials (bush – NCS DC 733348, chicken – NCS ZC73016 and pork muscle – NCS ZC 81001) for other elements.

Based on current understanding of the taxonomy and species interactions (Chen and Wise, 1999; Ponsard and Arditi, 2000; Scheu and Falca, 2000; Larochelle, 1990; El-Danasoury, 2016), invertebrates were grouped into the following categories: (1) herbivores feeding on living plant material (2) detritivores feeding on dead plant material in the litter layer, (3) omnivores that have variable diets (animals with admixture of plant material), and (4) predators with prey sources (Supplement 1); litter was assumed the basal resource of the detritus based food web.

## Statistical analysis

Differences between metal concentrations in litter and in invertebrates from various localities were compared using a one-way ANOVA with Tukey’s HSD post-hoc test. If the data did not meet normality and homogeneity of variance, we used nonparametric test (Kruskall-Wallis). To analyze the differences between trophic groups with regard to all analyzed elements, we performed a principal component analysis (PCA) on the correlation matrices.

All statistical analyses were performed using Statistica 10.0 (StatSoft Inc.). The stoichiometric ratios are reported as molar ratios.

## Results

### Elemental concentration in litter

Along the zinc pollution gradient in litter originating from the Olkusz sites the OH site contained significantly (F_2,10_=512, p<0.0001) more zinc (449 mg kg^−1^) than the other two sites with 158 and 147 mg Zn per kg litter for the OM and OR sites, respectively, that did not differ (Table 1). Copper concentrations in litter were significantly (F_2,10_=6.27, p<0.01) higher at the OM site (55.54 mg kg^−1^), but did not differ between OH and OR (20.4 and 8.19 mg k^−1^, respectively) (Table 1). The concentration gradient of iron followed that of zinc, however, with lower concentrations (F_2,12_=229, p<0.0001) (Supplement 2). Manganese had a counter gradient with highest concentrations at the reference site and lower ones at the polluted sites H_2,12_=12.5, p<0.01). The other elements (except Na) vary significantly among sites but with difference between all sites (K, Ca) or between OH and OR or OM (Mg, P) (Supplement 2). The significant differences in carbon and nitrogen concentrations were between OR and OM (C, F_2,12_=4.23, p<0.05) and between all sites (N, F_2,12_=184, p<0.0001).

At the Głogów transect litter concentrations of copper decreased with distance from the smelter (Table 1), however, only GH differed significantly from the other two sites (F_2,12_=7.29, p<0.05), with a concentration of 77.1 mg kg^−1^ the GM and GR site (24.8 and 13.9 mg kg^−1^ respectively) and the two last ones did not differ from one another. Zinc concentration in litter from the Głogów transect was significantly lower only in GM site than in GH and GR (F_2,12_=6.7, p<0.05). Insignificant were differences in litter concentration of C and Na. The other elements significantly varied between sites: GH from GM,GR (N, Ca) or GR from GM, GH (Mg, P, K, Ca, Mn) (Supplement 2).

### Elemental composition of trophic groups

Differences in elemental composition have been found between trophic groups at all the sites studied (Suppl. 2). The statistically significant differences were noted between litter and animals (especially predators) and among trophic groups – feeding on plant material (herbivores, detritivores) and those feeding mainly on other animals (omnivores, predators), but no particular pattern was observed (Suppl. 2). Significantly lower concentrations in litter than in animals were noted for N, Na, P, K and higher for Fe. The patterns for C, Mg, Cu and Zn were related to study site and transect.

The differences between herbivores, detritivores, omnivores and predators varied between particular elements and the sites studied (Suppl. 2). Looking at nitrogen, significantly higher was the concentration of this element in omnivores than herbivores and detritivores (sites OM- F_3,44_=12.8, p<0.001 and OR – (F_3,66_=7.6, p<0.001), but in OH and Głogów transects no significant differences were found. In phosphorus concentration the significant difference was found only in OR – detritivores were richer than predators (F_3,66_=6.3, p<0.001). Clear pattern was observed in sodium concentration: litter < herbivores < detritivores < omnivores < predators. Significant differences between consumers were found: herbivores and omnivores/predators (OH – F_3,66_=55.3, p<0.0001, OM – F_3,66_=3.5, p<0.05), detritivores/herbivores and predators, herbivores and omnivores (OR – F_3,67_=32.7, p<0.0001), herbivores and omnivores/predators (GM – F_3,66_=8.8, p<0.001), (Suppl. 2)

The PCA in some cases separated trophic groups (Fig. 1) and first and second axes explain 61 (OR, OM), 60 (OH, GR), 67 (GM) and 77 (GH) percent of variance. In all sites studied litter samples create a group separate from consumers. In OR first axis and in OM second axis separate clearly herbivores and detritivores from omnivores and predators. However it is clear that composition of slugs, Isopods and Diplopods differs from those of beetles and spiders or ants. At all studied sites the positions of herbivorous species like *Arion fuscus* (slug) and both *Amara aenaea* and *Hylobius abietis* (beetles) are placed separately on graph. Stoichiometry of trophic groups

**Fig. 1.**
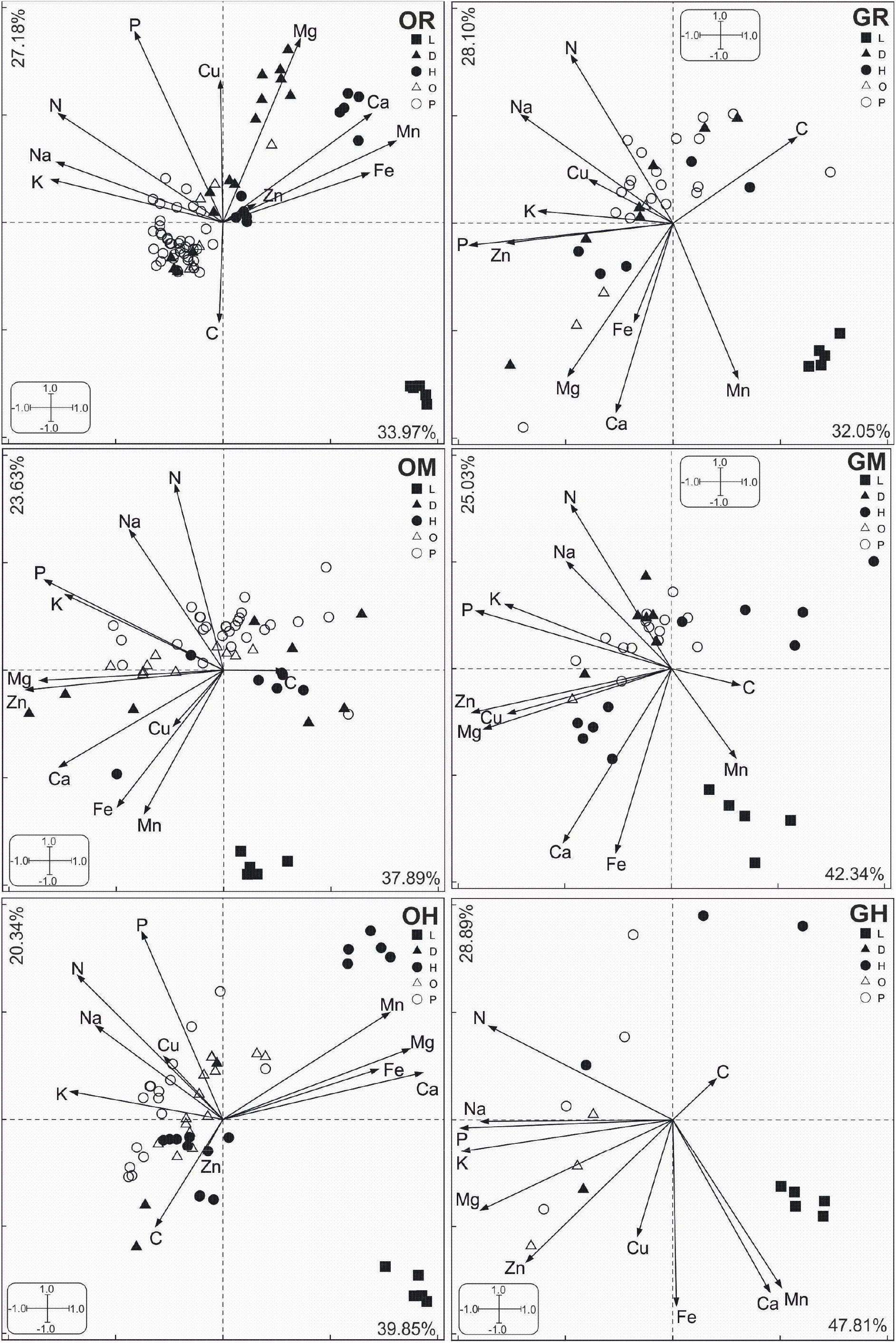
PCA ordination diagrams showing the relation of: the litter (L), herbivores (H), detritivores (D), omnivores (O) and predators (P) towards one another based on their elemental concentration. The six plots represent the Legnica sites (right) and Olkusz sites (left) where: GH – Głogów heavily polluted site, GM – Głogów moderately polluted site, GR – Głogów reference site; OH – Olkusz heavily polluted site, OM – Olkusz moderately polluted site, OR – Olkusz reference site.

At all six study sites element ratios in litter were significantly distinct from these in invertebrates with a higher C:N, C:Na, C:P, C:K ratio and lower C:Mg, C:Ca and C:Fe ratios. The N:P ratio show some differences between trophic groups achieving the highest values for detritivores (OH, OM) or the highest for herbivores (GH, GM), however the differences between groups were not significant. We calculated the resouce:consumer ratios for the N:P to check if studied trophic groups on transects are N limited or P limited. The ratio was calculated for litter as a food source for detritivores and herbivores, and for detritivores and herbivores as a food source for predators. Almost all values were below 1, only in some cases above 1, but the results were insignificant. It suggests that the trophic groups are rather N limited and not P limited.

### Comparison of trophic groups between sites on transects

Elemental composition of particular trophic groups was compared on transects Olkusz transect (between OH, OM, OR) and Głogów transect (between GH, GM, GR). Multivariate ANOVA shows that in Olkusz transect significant differences in elemental composition were both between sites (F_22_=194, p<0.0001) and between trophic groups (F_44_=373, p<0.0001). On Olkusz transect herbivores differed significantly in the concentrations of Na, Mg, P, K, Ca, Mn and Fe, however for Mn a significant difference was between OH and OR only, and in other elements significant differences were between OH, OR and OM. Significant differences in concentrations of Zn an Cu were observed in Olkusz transect only in predators: Zn – OH from OM, OR (F_3,59_=11.7, p<0.0001), Cu – OH from OR (F_3,59_=5.7, p<0.01).In Głogów transect Multivariate ANOVA shows that significant differences in elemental composition were both between sites (F_22_=94, p<0.01) and between trophic groups (F_44_=188, p<0.0001). However looking at particular elements within trophic groups the significant differences were found only in Cu concentration between omnivores from GM and GR (F_2,3_=14.4, p<0.005) and in Mn between predators from GM and both GH,GR (F_2,30_=7.97, p<0.05).

Compared were C:N and N:P ratios for particular trophic groups along pollution gradients (Olkusz Forest and Głogów Forest), but obtained results did not bring conclusive findings. In herbivores statistically significant difference were found in C:Zn ratio (between OH and OM, F_2,19_=6.70, p<0.05). Comparing only C:Zn between OH and OR there was significant difference (F_1,33_=16.0, p<0.00) with higher ratio in OR. Detritivores differed in ratios C:Cu (F_2,50_=3.2, p<0.05, OM from OH,OR) and C:Zn (F_2,59_=4.7, p<0.05, OM from OH,OR). On the Głogów transect no significant differences between sites for particular trophic groups have been found.

### Trophic groups in beetles

Taking into account the differences in body composition of particular taxa creating one trophic group, only Coleoptera were considered. Results of PCA (Fig. 2) show that there are differences in body composition between species within one trophic group. It is clearly visible on graphs for OH and GM where two herbivores *A. aenaea* and *H. abietis* are separated one form another, and *C. nemoralis* segregates from other predatory beetles (OH).

**Fig. 2.**
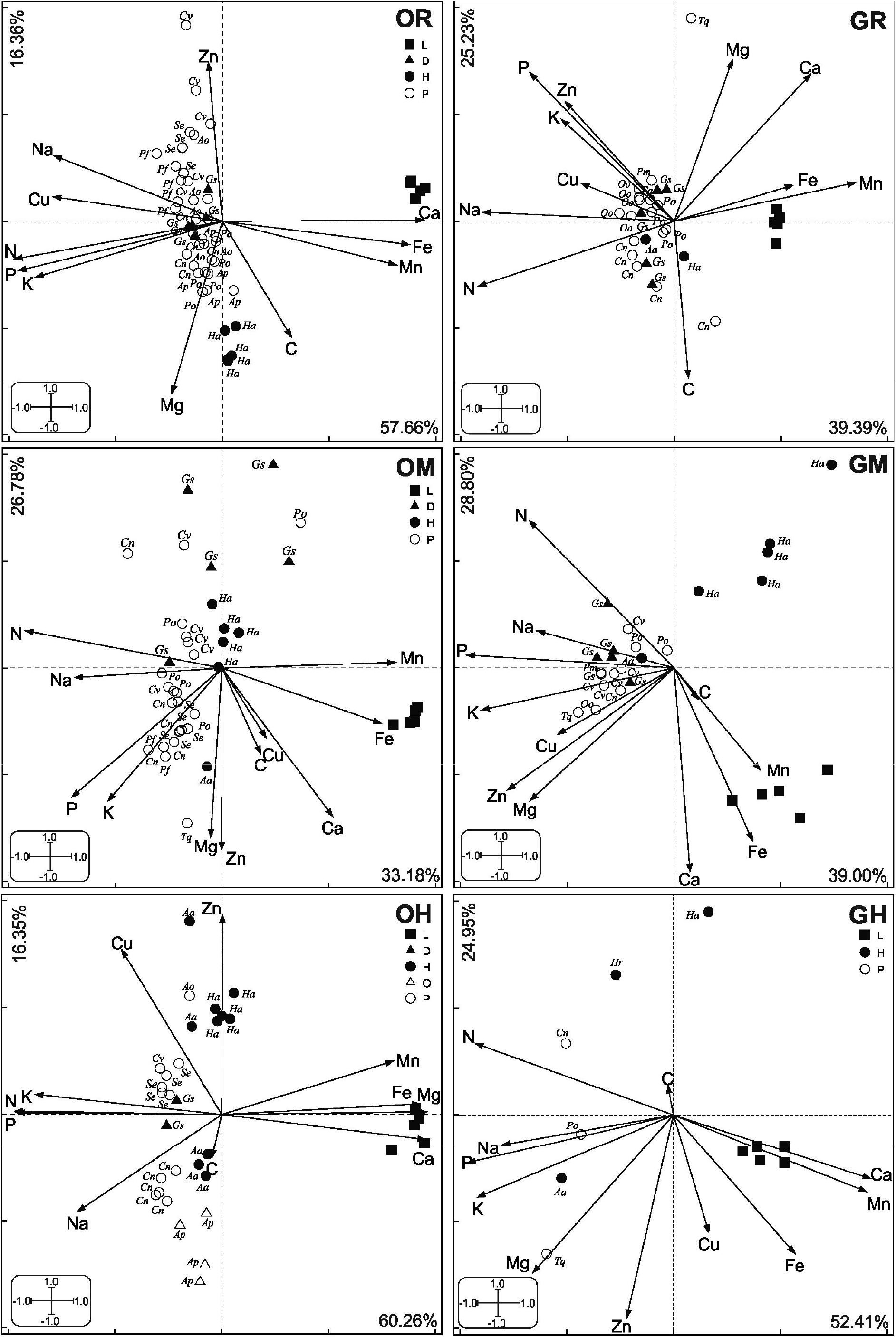
PCA ordination diagrams for Coleoptera showing the relation of: the litter (L), herbivores (H), detritivores (D), omnivores (O) and predators (P) towards one another based on their elemental concentration. The six plots represent the Legnica sites (right) and Olkusz sites (left) where: GH – Głogów heavily polluted site, GM – Głogów moderately polluted site, GR – Głogów reference site; OH – Olkusz heavily polluted site, OM – Olkusz moderately polluted site, OR – Olkusz reference site. Species abbreviation in Supplement 1.

### Faunal composition and abundance

The studied transects differed in taxonomic richness, diversity and abundance of litter dwelling invertebrates. Generally, the Olkusz sites were more densely populated than the Głogów sites; at the Olkusz sites on average 2992 individuals per site were sampled, whereas at the Legnica sites only 555 individuals per site on average were captured. Corresponding to the higher abundance, the taxonomic richness found at the Olkusz sites was higher than that at the Głogów sites (averages of 19 and 14 taxonomic groups per site, respectively). Most abundant at almost all sites were Formicidae accounting for more than 60% of the individuals. Only in OR dominant were Carabidae.

## Discussion

In the present study litter on polluted sites was enriched with Zn or Cu as a result of industrial pollution in doses which cause the concentrations exceeding maximum permissible concentrations in soil (40 mg×kg^−^1 for Cu and 160 mg×kg^−1^ for Zn, Crommentuijn et al., 2000), what was expected to affect body composition and stoichiometry of consumers living there. Observed were significant differences in both pollutants concentration in litter on transects both in Olkusz Forest (Zn) and Głogów Forest (Cu). In the studied transects the sites differed not only in concentrations of Zn and Cu in litter, but in the concentrations of other elements as well, as it has been shown in the analysis performed. It is an unavoidable problem of all field studies because a uniform quality and composition of litter is may be available possible only in laboratory experiments. In the field always numerous factors cause differences between sites.

Dead organic matter on the forest floor is consumed by detritivores and omnivores (feeding on mixed animal and plant material) and these both groups as well as herbivores constitute the prey of predators (Chen and Wise, 1999; Ponsard and Arditi, 2000; Scheu and Falca, 2000). The litter quality and environmental conditions affect density and diversity of litter organisms (Dyer and Letourneau, 2003), as well as C:X ratios and interaction between elements in the trophic web (Ott et all., 2014). The higher abundance and diversity of litter fauna in Olkusz Forest than in Głogów Forest was probably the result of environmental conditions: lower precipitation in Głogów region as well as a thinner litter layer what creates inconvenient conditions for the organisms living there.

The body tissues of organisms are built of approximately 25 chemical elements (Sterner and Elser, 2002; Kaspari et al., 2016) and tissues of plants and animals differ in content of proteins, carbohydrates and lipids. The differences observed are similar in elemental composition and stoichiometry. Literature data suggest that animals feeding on food poor in some elements will have lower concentration of these elements than those feeding on a richer food. Therefore herbivores and detritivores should have lower concentration of N and P in their bodies than predatory species (Elser et al., 2000; Fagan et al., 2002; Fagan and Denno, 2004; Feijoó et al., 2014; Gonzalez et al., 2011; Lemoine et al., 2014). Similarly to the data cited above in Olkusz Forest the concentrations of N were higher in predatory and omnivorous taxa than in first order consumers and detritus. However our results for P do not confirm observations of other authors. The highest concentration of P was found in various trophic groups in studied sites. A possible explanation is that the differences exist in taxonomical composition of trophic groups in studied sites. The herbivores included beetles (A. *aenea, H. abietis)* as well as Gastropoda (*Ar.fuscus*). The group of detritivores included beetles, Isopoda and Diplopoda, the omnivores were beetles, harvestman and ants while the predator group was composed of beetles, spiders and chilopods. The taxa such as gastropods, chilopods and especially isopods and diplopods contain high concentrations of P and Ca. Therefore its share in trophic group may have affected pattern of trophic group differences in the transect studied. Impact may have additionally differentiation in body size of animals within trophic groups, since the literature data bring information on negative allometry between P and body mass (Woods et al. 2004, Hambäck et al. 2009). Because of the above results we decided to limit taxonomical diversity and only Coleoptera were taken into analyses. Our results of PCA (Fig 2) show that still some herbivorous beetles (*A.aenea* and *H. abietis*) are differently positioned in the multidimensional space, as shown on the graph (Fig. 2). The results seem to support the thesis on importance of taxonomical identity over trophic group (Gonzalez et al., 2011, 2018). As it has been highlighted by Gonzalez et al (2011) the phylogeny may exert significant influence on the results for trophic groups. Taxonomy explains most of the variance in elemental composition, and there is strong dependency of macroinvertebrate stoichiometry on taxonomy and trophic group (Gonzalez et al 2018).

The topic of interest in our study was if metal pollution affect ratios among elements e.g. C:N or N:P. According to Sardans et al. (2012) the ratios between two main elements, carbon and nitrogen, differ between particular trophic groups: C:N_plants_ >> C:N_herbivores_ > C:N_predators_. In our results the changes in the C:N ratios were decreasing as trophic level increased, the C:N ratio in litter was 10 fold higher than in detritivores. Similarly, Bradshaw et al. (2012) found that within one trophic link all interactions have similar elemental patterns. In our study the C:N ratio in predators was similar to (or lower than) in herbivores, because this animals feed on source reach in necessary elements (Gonzalez et al., 2018). Detritivores are feeding on food with C:N ratio ten times higher than its body, and higher C:Na, C:P, C:K ratios in litter, what suggest that these animals can be confronted with “stoichiometric mismatch” and their food should be supplemented from other sources (Filipiak and Weiner, 2014, 2016). Therefore detritivores as well as herbivores complement elements in shortage in their food by overfeeding or compensatory feeding (Sterner and Elser, 2002). We hypothesized that animals living in metals polluted sites will have lower C:N ratio because C is related with lipid storage, and costs of detoxification will result in smaller fat reserves. However such relation was not found. With similar observation yielded study on *P. oblongopunctatus* in Olkusz transect. Zygmunt et al. (2006) did not found significant trend in body caloric value on pollution transect.

The N:P ratio did not differ either between trophic groups or study sites. The calculated resource:consumer N:P ratio suggests that invertebrates in the forests studied are N limited and not P limited what is consistent with other data (Lemoine et al., 2014).

To our knowledge this is the first study that describes the elemental differences between trophic groups in a terrestrial detritivore food web both with and without the presence of metal pollution. As expected, litter was poor in elements such as nitrogen and phosphorus.

Further, similar element ratios of the different trophic groups of invertebrates at the different sites suggests that metal pollution has little effect on the overall elemental balance in terrestrial detritivorous macro-invertebrate food webs.

## Supporting information

Species list of selected taxa with trophic classification

Averages and standard deviations of elemental concentrations

## Acknowledgements

The authors would like to thank the Polish Foundation for Sciences (grant: MPD/2009-3/5/) for their financial support and the Polish National Science Centre (grant: 2012/05/N/NZ8/00985).

