## Supplementary material for "Does metal pollution affect stoichiometry of soil-litter food webs?": Species list of selected taxa with trophic classification

**Supplement 1:** Species list of selected taxa with trophic classification (TL = Trophic level; SA = species abbreviation; Herb. = herbivores; Detr. = detritivores; Omni. = omnivores; Pred. = predators) and number of individuals per site trapped and used for analysis. The “X” in the row with Collembola indicates presence but no numbers counted.

| | | | Number of individuals per site | | | | | |
|---|---|---|---|---|---|---|---|---|
| **Taxa** | **SA** | **TL** | **LH** | **LM** | **LR** | **OH** | **OM** | **OR** |
| *Amara aenea* | Aa | Herb. | 3 | 17 | 3 | 45 | 9 | 2 |
| *Arion fuscus* | Af | Herb. | | 105 | 13 | 73 | 6 | 27 |
| *Hylobius abietis* | Ha | Herb. | 4 | 24 | 6 | 27 | 348 | 41 |
| Collembola | | Detr. | X | X | X | X | X | X |
| Diplopoda | | Detr. | | | | | 8 | 47 |
| *Geotrupes stercorosus* | Gs | Detr. | | 27 | 31 | 2 | 13 | 286 |
| Isopoda | | Detr. | | | 2 | | | 6 |
| *Harpalus rufipes* | Hr | Omni. | 4 | | | 3 | | 4 |
| *Abax parallepipedus* | Ap | Omni. | 1 | | | 8 | 8 | 12 |
| *Formicidae* | | Omni. | 139 | 953 | 89 | 1682 | 3218 | 550 |
| Opilionidae | | Omni. | 1 | 12 | 7 | 14 | 38 | 15 |
| *Abax ovalis* | Ao | Pred. | | 1 | 7 | 3 | 3 | 86 |
| *Carabus violaceus* | Cv | Pred. | | 7 | | 1 | 15 | 140 |
| *Lithobius spp.* | | Pred. | | 1 | | 25 | 23 | 72 |
| *Carabus nemoralis* | Cn | Pred. | 1 | 2 | | 131 | 13 | 215 |
| *Ocypus olens* | Oo | Pred. | | 1 | 13 | 151 | 25 | 14 |
| *Parabemus fossor* | Pf | Pred. | | | | | 18 | 24 |
| *Pterostichus metallicus* | Pm | Pred. | | 5 | 2 | | | |
| *Pterostichus oblopunctatus* | Po | Pred. | 2 | 3 | 13 | | 33 | 185 |
| *Staphylinus erythropterus* | Se | Pred. | | | | 844 | 191 | 125 |
| *Tachys quadrisignatus* | Tq | Pred. | 14 | 24 | 9 | | 7 | |
| Thomisidae | | Pred. | | 33 | 12 | 11 | 14 | 11 |
| Other spiders | | Pred. | 53 | 14 | 5 | 11 | 84 | 8 |
