## Supplementary material for "Does metal pollution affect stoichiometry of soil-litter food webs?": Averages and standard deviations of elemental concentrations

**Supplement 2:** Averages and standard deviations of elemental concentrations in litter and the trophic groups at all six sites.

| Site | Group | | C % | N % | Na mg/kg | Mg mg/kg | P mg/kg | K mg/kg | Ca mg/kg | Mn mg/kg | Fe mg/kg | Cu mg/kg | Zn mg/kg |
|---|---|---|---|---|---|---|---|---|---|---|---|---|---|
| OH | Litter | Mean | 47.6 | 1.0 | 367.4 | 2056.2 | 781.5 | 3016.5 | 10261.4 | 83.4 | 580.3 | 20.4 | 448.7 |
| | | SD | 0.4 | 0.1 | 274.9 | 107.6 | 41.5 | 246.5 | 624.9 | 5.6 | 17.6 | 7.8 | 15.3 |
| OH | Herbivores | Mean | 47.0 | 9.6 | 648.4 | 1588.3 | 8062.4 | 4477.9 | 4377.6 | 143.2 | 341.6 | 64.5 | 1096.4 |
| | | SD | 3.2 | 1.4 | 324.9 | 999.1 | 4287.4 | 1097.9 | 5698.3 | 183.7 | 316.1 | 36.1 | 1621.8 |
| OH | Detritivores | Mean | 50.8 | 9.6 | 1206.3 | 736.5 | 5071.2 | 4980.0 | 932.0 | 19.2 | 348.8 | | |
| | | SD | 3.5 | 2.3 | 530.8 | 155.2 | 1335.7 | 917.7 | 925.3 | 19.4 | 450.8 | | |
| OH | Omnivores | Mean | 48.4 | 10.4 | 2115.5 | 978.0 | 6456.6 | 5545.4 | 781.5 | 30.5 | 328.0 | 83.4 | 580.3 |
| | | SD | 1.4 | 0.6 | 361.6 | 302.2 | 1560.7 | 992.9 | 1264.8 | 32.6 | 153.4 | 11.5 | 178.3 |
| OH | Predators | Mean | 46.9 | 10.2 | 2484.1 | 972.1 | 6651.3 | 5589.3 | 1199.7 | 25.6 | 126.8 | 83.2 | 494.3 |
| | | SD | 3.7 | 1.1 | 522.2 | 441.3 | 1718.7 | 1420.2 | 1242.7 | 22.8 | 117.3 | 99.2 | 268.0 |
| OM | Litter | Mean | 53.5 | 1.8 | 59.0 | 522.0 | 869.1 | 1180.6 | 4598.6 | 240.2 | 248.6 | 55.5 | 157.8 |
| | | SD | 5.6 | 0.1 | 30.1 | 84.7 | 118.3 | 273.5 | 571.7 | 35.8 | 37.7 | 33.8 | 19.2 |
| OM | Herbivores | Mean | 48.3 | 9.2 | 365.2 | 605.6 | 3676.0 | 2037.3 | 1283.4 | 460.7 | 89.1 | 92.6 | 191.9 |
| | | SD | 4.7 | 1.2 | 696.8 | 226.0 | 2352.6 | 419.1 | 2038.6 | 924.5 | 66.4 | 97.6 | 263.3 |
| OM | Detritivores | Mean | 44.0 | 9.8 | 803.1 | 833.3 | 5668.2 | 2910.4 | 21048.1 | 128.2 | 208.4 | 55.1 | 275.2 |
| | | SD | 5.7 | 1.1 | 259.8 | 987.5 | 7421.4 | 2789.8 | 43605.9 | 116.3 | 244.4 | 55.2 | 320.0 |
| OM | Omnivores | Mean | 52.2 | 12.5 | 974.2 | 801.2 | 5623.1 | 3069.5 | 1351.9 | 153.0 | 169.1 | 31.2 | 303.7 |
| | | SD | 5.5 | 1.2 | 375.7 | 358.5 | 2248.5 | 900.4 | 796.5 | 76.7 | 46.5 | 37.6 | 193.7 |
| OM | Predators | Mean | 51.6 | 11.2 | 900.2 | 676.0 | 4419.5 | 2572.3 | 774.6 | 56.2 | 64.6 | 33.6 | 220.5 |
| | | SD | 3.3 | 1.3 | 387.5 | 405.3 | 1930.9 | 1094.6 | 664.5 | 42.6 | 49.1 | 31.3 | 174.1 |
| OR | Litter | Mean | 49.4 | 0.8 | 180.9 | 635.4 | 611.4 | 2046.3 | 9078.6 | 452.2 | 311.9 | 8.2 | 147.2 |
| | | SD | 0.4 | 0.0 | 55.0 | 18.2 | 10.2 | 100.5 | 179.7 | 8.7 | 17.2 | 1.1 | 2.9 |
| OR | Herbivores | Mean | 45.9 | 8.6 | 413.2 | 1550.4 | 7314.6 | 4018.9 | 4325.8 | 2308.9 | 216.1 | 86.4 | 443.4 |
| | | SD | 2.7 | 1.0 | 199.3 | 737.7 | 2486.7 | 1869.0 | 3895.4 | 2609.0 | 144.4 | 34.5 | 485.4 |
| OR | Detritivores | Mean | 39.5 | 8.3 | 1689.7 | 1819.8 | 10010.6 | 4743.2 | 42297.9 | 166.8 | 173.6 | 38.0 | 104.9 |
| | | SD | 5.6 | 2.2 | 689.6 | 1248.8 | 5213.0 | 602.9 | 47007.8 | 162.3 | 133.0 | 22.8 | 29.7 |
| OR | Omnivores | Mean | 47.2 | 11.2 | 1954.5 | 927.8 | 6865.6 | 5905.2 | 2151.8 | 366.1 | 209.6 | 20.0 | 149.3 |

| | | | | | | | | | | | | | |
|---|---|---|---|---|---|---|---|---|---|---|---|---|---|
| | | SD | 1.6 | 0.7 | 443.1 | 431.8 | 2933.9 | 1230.9 | 2669.1 | 593.9 | 108.9 | 3.5 | 10.0 |
| OR | Predators | Mean | 46.9 | 9.4 | 2312.9 | 801.4 | 6205.4 | 5571.8 | 842.8 | 53.8 | 73.2 | 27.1 | 168.2 |
| | | SD | 4.9 | 1.1 | 559.1 | 187.5 | 1487.9 | 1357.6 | 698.2 | 32.7 | 30.8 | 9.3 | 102.1 |

4 **Supplement 2 Continued**

| Site | Group | | C % | N % | Na mg/kg | Mg mg/kg | P mg/kg | K mg/kg | Ca mg/kg | Mn mg/kg | Fe mg/kg | Cu mg/kg | Zn mg/kg |
|---|---|---|---|---|---|---|---|---|---|---|---|---|---|
| GH | Litter | Mean | 51.6 | 1.1 | 104.3 | 226.3 | 408.4 | 417.0 | 4479.6 | 757.2 | 192.9 | 77.0 | 70.4 |
| | | SD | 1.1 | 0.09 | 110.9 | 23.0 | 25.1 | 69.9 | 388.1 | 79.6 | 14.4 | 32.5 | 4.8 |
| GH | Herbivores | Mean | 48.4 | 8.9 | 318.1 | 272.8 | 2015.2 | 1041.9 | 768.8 | 61.7 | 1.3 | 94.1 | 47.8 |
| | | SD | 10.3 | 0.89 | 300.1 | 210.1 | 1858.2 | 996.0 | | | | 121.2 | 56.4 |
| GH | Detritivores | Mean | 54.4 | 15.0 | 1106.6 | 725.1 | 7217.6 | 4183.5 | 2348.5 | 335.7 | 342.0 | 94.1 | 113.4 |
| | | SD | 0.0 | 0.00 | | | | | | | | | |
| GH | Omnivores | Mean | 46.8 | 11.3 | 1324.4 | 896.1 | 6760.4 | 3541.8 | 1934.5 | 263.8 | 120.1 | 71.5 | 307.6 |
| | | SD | 1.9 | 2.19 | 734.6 | 434.6 | 3856.5 | 928.1 | 1872.1 | 61.5 | 52.4 | 44.7 | 230.6 |
| GH | Predators | Mean | 54.8 | 10.6 | 747.1 | 691.6 | 4241.1 | 2453.0 | 993.3 | 49.3 | 71.2 | 110.0 | 180.2 |
| | | SD | 5.2 | 1.38 | 418.4 | 413.2 | 2037.6 | 1110.2 | 1093.6 | 26.2 | 56.1 | 89.6 | 146.1 |
| GM | Litter | Mean | 49.3 | 1.1 | 293.1 | 331.2 | 391.9 | 570.0 | 6301.8 | 742.1 | 250.0 | 24.8 | 50.6 |
| | | SD | 1.30 | 0.2 | 365.5 | 19.1 | 55.0 | 297.5 | 603.5 | 69.6 | 73.3 | 16.2 | 11.1 |
| GM | Herbivores | Mean | 48.2 | 9.4 | 351.9 | 614.3 | 3657.0 | 770.8 | 4499.5 | 276.8 | 128.1 | 112.8 | 136.8 |
| | | SD | 3.44 | 0.9 | 250.8 | 548.8 | 2866.7 | 487.0 | 4705.7 | 472.0 | 74.1 | 112.0 | 132.8 |
| GM | Detritivores | Mean | 43.9 | 10.5 | 666.5 | 587.1 | 3473.8 | 2952.4 | 906.2 | 41.9 | 82.9 | 47.5 | 85.1 |
| | | SD | 6.38 | 1.6 | 335.4 | 144.8 | 1886.5 | 880.9 | 1067.2 | 38.5 | 70.9 | 37.0 | 32.4 |
| GM | Omnivores | Mean | 49.0 | 12.2 | 1524.4 | 1234.1 | 7628.1 | 4701.4 | 3076.8 | 1055.1 | 181.8 | 182.2 | 287.5 |
| | | SD | 0.00 | 0.0 | | | | | | | | | |
| GM | Predators | Mean | 48.9 | 10.5 | 749.2 | 594.0 | 3854.7 | 2220.8 | 609.1 | 315.2 | 43.7 | 66.0 | 174.6 |
| | | SD | 6.40 | 1.0 | 323.5 | 259.8 | 1192.3 | 796.9 | 330.2 | 222.3 | 27.8 | 67.6 | 128.3 |
| GR | Litter | Mean | 55.4 | 2.4 | 49.9 | 683.9 | 1283.5 | 1478.3 | 5440.6 | 1011.4 | 93.6 | 13.9 | 66.4 |
| | | SD | 6.4 | 0.2 | | 55.2 | 87.5 | 95.7 | 587.5 | 120.0 | 7.0 | 18.4 | 10.0 |
| GR | Herbivores | Mean | 48.9 | 10.2 | 406.4 | 790.7 | 5979.4 | 1818.8 | 5506.5 | 90.7 | 102.2 | 78.8 | 208.5 |
| | | SD | 2.1 | 0.6 | 242.5 | 291.8 | 2701.5 | 387.6 | 4778.5 | 71.0 | 64.7 | 47.0 | 131.1 |
| GR | Detritivores | Mean | 49.2 | 10.9 | 1071.4 | 883.1 | 4950.8 | 3642.7 | 12263.1 | 160.7 | 101.0 | 154.3 | 119.3 |
| | | SD | 10.9 | 3.2 | 753.1 | 621.6 | 2686.9 | 1475.3 | 30054.3 | 199.4 | 54.9 | 215.9 | 61.7 |
| GR | Omnivores | Mean | 49.0 | 11.9 | 1481.4 | 1108.7 | 6735.3 | 4298.1 | 1865.0 | 604.3 | 2009.2 | 10.6 | 213.9 |

| | | | | | | | | | | | | | |
|---|---|---|---|---|---|---|---|---|---|---|---|---|---|
| | | SD | 3.3 | 1.0 | 537.5 | 632.3 | 2461.4 | 828.9 | 1230.7 | 359.7 | 3113.1 | 13.3 | 101.2 |
| GR | Predators | Mean | 51.7 | 10.6 | 1072.4 | 585.6 | 3772.5 | 2409.4 | 534.2 | 135.1 | 50.5 | 44.4 | 171.4 |
| | | SD | 3.8 | 1.3 | 555.2 | 249.2 | 1463.1 | 1136.3 | 283.8 | 109.5 | 25.9 | 29.7 | 111.1 |
